# Regular physical activity reduces the percentage of spinally projecting neurons that express mu-opioid receptors from the RVM

**DOI:** 10.1101/2020.04.27.064923

**Authors:** KA Sluka, SJ Kolker, J Danielson, L Rasmussen

## Abstract

Regular physical activity/exercise is an effective non-pharmacological treatment for individuals with chronic pain. Central inhibitory mechanisms, involving serotonin and opioids, are critical to analgesia produced by regular physical activity. The RVM sends projections to the spinal cord to inhibit or facilitate nociceptive neurons and plays a key role in exercise-induced analgesia. The goal of these studies was to examine if regular physical activity modifies RVM-spinal cord circuitry. Male and female mice received Fluoro-Gold placed on the spinal cord to identify spinally projecting neurons from the rostral ventromedial medulla (RVM) and the nucleus raphe obscuris/nucleus raphe pallidus (NRO/NRP), dermorphin-488 into caudal medulla to identify mu-opioid receptors, and were immunohistochemically stained for either phosphorylated-N-methyl-D-aspartate subunit NR1 (p-NR1) to identify excitatory neurons or tryptophan hydroxylase (TPH) to identify serotonin neurons. The percentage of dermorphin-488-positive cells that stained for p-NR1 (or TPH), and the percentage of dermorphin-488-positive cells that stained for p-NR1 (or TPH) and Fluoro-Gold was calculated.

Physically active animals were provided running wheels in their cages for 8 weeks and compared to sedentary animals without running wheels. Animals with chronic muscle pain, induced by two intramuscular injections of pH 4.0, were compared to sham controls (pH 7.2). Physically active animals had less mu-opioid expressing neurons projecting to the spinal cord when compared to sedentary animals in the RVM, but not the NRO/NRP. No changes were observed for TPH. These data suggest that regular exercise alters central facilitation so that there is less descending facilitation to result in a net increase in inhibition.

**Summary Statement:** Physically active animals has less mu-opioid expressing neurons projecting to the spinal cord in the RVM, but not the NRO/NRP, when compared to sedentary animals.

## Introduction

Regular physical activity/exercise is an important and effective non-pharmacological treatment for individuals with chronic pain [3,8-10,26], and large population studies show that people who are physically active have a lower incidence of chronic pain [34,35,69]. In parallel, prior studies show that regular physical activity prevents development of chronic muscle pain, activity-induced pain, and neuropathic pain in animal models [4-6,19,37,39,58]. Central inhibitory mechanisms, involving serotonin and opioids, are critical to the analgesia produced by regular physical activity. Specifically, there are increases in endogenous opioids in the rostral ventromedial medulla (RVM) and the periaqueuctal gray (PAG) with regular exercise, and blockade of opioid receptors systemically or supraspinally (RVM, PAG) reduces the analgesic effects of regular physical activity in animal models of chronic muscle pain and neuropathic pain [6,39,60]. Further, regular physical activity increases serotonin and decreases the serotonin transporter while systemic depletion of serotonin prevents the analgesia in animal models of chronic muscle pain and neuropathic pain [4,6,39]. These data show that central inhibitory pathways, including the RVM, are important components of exercise-induced analgesia.

Prior data show that there is an increase in phosphorylated-N-methyl-D-aspartate subunit p-NR1 (ser 831) (p-NR1) in the RVM and caudal medulla (NRO/NRP) in response to repeated injections of acidic saline in sedentary mice and that regular physical activity prevents this increase [58]. The RVM contains 3 populations of neurons: ON cells, OFF cells, and neutral cells [16]. The ON cells are facilitation neurons that when activated enhance pain. Experimentally, ON cells express mu-opioid receptors (MOR), are directly inhibited by MOR agonists, and removal of MOR-expressing neurons in the RVM with dermorphin-saporin prevents the development of hyperalgesia to nerve injury [16,25,30,33,48,49]. The RVM and caudal medulla also contain serotonergic cells, proposed to be neutral cells [50,51]. However, increases in serotonin in the RVM produces analgesia, and blockade of serotonin receptors or the serotonin transporter in sedentary animals are analgesic [22,27]. Further, there are interactions between the serotonin and opioid systems within the RVM [18]. Specific to exercise-induced analgesia blockade of opioid receptors prevents the exercise-induced reductions in the serotonin transporter [39]. Thus, ON-cells, MOR, and serotonin in the RVM are key components of endogenous inhibition.

The RVM sends projections to the spinal cord to inhibit or facilitate nociceptive neurons in the dorsal horn. Prior studies show that nearly 60% of spinally projecting RVM neurons respond to mu-opioid agonists, 40% of spinally projecting neurons express the serotonin marker tryptophan hydroxylase, and a subpopulation of spinally projecting neurons expression mu-opioid receptors [41]. However, it is unclear how exercise modulates the circuitry in the RVM. The goal of these studies was to examine the ability of regular physical activity to modify the RVM-spinal cord circuitry. We hypothesized that there were would be a reduction in spinally projecting mu-opioid expressing neurons to the spinal cord, a reduction in p-NR1 in mu-opioid-expressing neurons, and no change in trypotophan hydroxylase-projecting spinal neurons.

## Materials & Methods

All animal procedures were approved by the authors’ institution’s animal care and use committee. C57BL/6 mice (Jackson Laboratories) were used for these experiments. For dermorphin-488 analysis, Male (n=10) and female (n=11) C57/Black 6 mice (Jackson Laboratories) were used for these studies. Mice received either repeated injections of pH 4.0 saline (n=14) or pH 7.2 saline (n=7) and were either sedentary (n=11) or physically active (n=10). A subpopulation of animals was analyzed for TPH. Male (n=7) and female (n=8) mice were used for these studies. Mice received either repeated injections of pH 4.0 saline (n=8) or pH 7.2 saline (n=7), and were either sedentary (n=8) or physically active (n=7). Physically active mice were housed individually with running wheels in their cages for 8 weeks while sedentary mice were housed individually in home-cages without running wheels. Prior studies show that there were no sex differences in the analgesia [37]. Two 20 μl injections of saline (pH 4.0 or pH 7.2) were given into the gastrocnemius muscle, 5 days apart, while the animal was anesthetized with 2-4% isoflurane [57]. Fluoro-Gold was applied to the spinal cord 7 days before perfusion to label spinally projecting neurons from the rostrocaudal medulla. Dermorphin-488 was injected 60 minutes before perfusion 24h after the second intramuscular saline injection to label mu-opioid receptors.

### Labeling of spinally projecting neurons

To label spinally projecting neurons, Fluoro-Gold was applied to the spinal cord. Mice were anesthetized with isoflurane (2-5%) and a laminectomy was performed to expose the lumbar spinal cord (L4-L6 region). A Fluoro-Gold (2%) soaked piece of gel foam was applied to the surface of the L4-5 dorsal horn and left in place. Animals were sutured closed and allowed to recover for 2 days prior to intramuscular saline injections.

### Labelling cells with mu-opioid receptors

Preliminary experiments in mice were unable to find a mu-opioid receptor antibody that showed staining selectivity for wild-type mice but not mu-opioid mice in the rostrocaudal medulla (RVM). We therefore directly injected dermorphin conjugated to a Hilyte-488 fluorescent tag into the RVM to bind and label cells expressing mu-opioid receptors. Dermorphin-488 is internalized after binding to MOR and thus represents cells that express mu-opioid receptors[1,40]. Control experiments showed no dermorphin-488 signal in mu-opioid receptor knockout mice (see Figure 1A, B).

**Figure 1.**
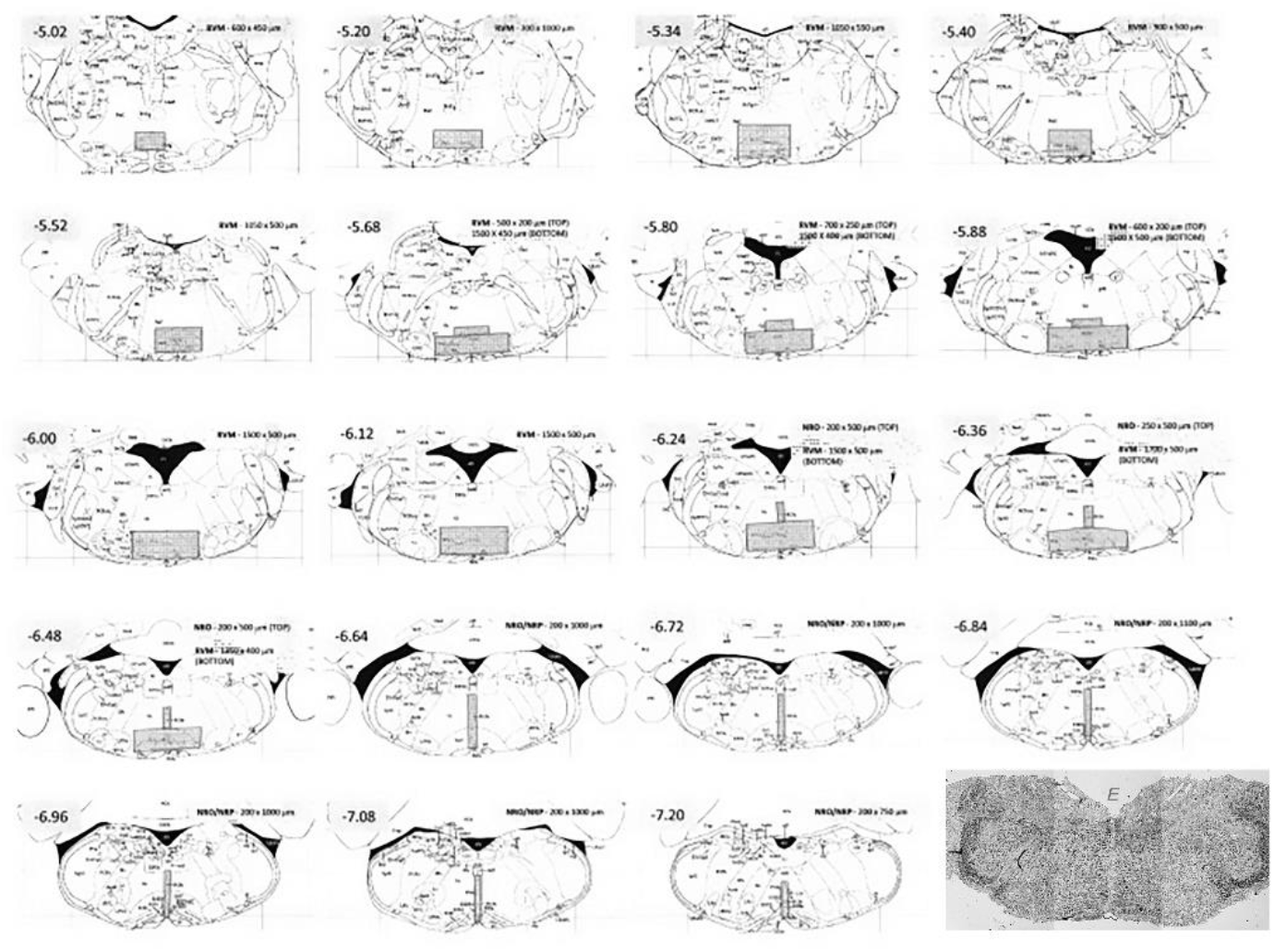
Figure 1 shows a template used to identify areas of the RVM, NRO or NRP by Bregma defined by Paxinos and Franklin’s Mouse Atlas [46]. Shaded areas show the identified areas quantified for the RVM, NRO and NRP. A light level phase image is also shown. Light level images of the medulla were used to identify the location of the section analyzed and identify areas for confocal imaging at higher magnification.

Mice were anesthetized with Ketamine/Xylazine (6μl/g) and were placed into stereotaxic frame. Multiple injections of dermorphin-488 (0.2 mg/ml in 20% DMSO, 0.2 μl) were made into the RVM, NRO and NRP. A 33-gauge injector was attached to a piece PE10 tubing and a Hamilton syringe. The Hamilton syringe and tubing were filled with saline, then fluid was extruded from the tubing, a bubble was introduced into the tubing and then 1-2 μl of dermorphin was drawn up to fill the tubing. Prior to each injection a small amount of dermorphin was injected out of the end of the needle to ensure the tip was filled. Three injections of dermorphin-488 were injected directly in the rostrocaudal medulla 0.1 mm in-between each injection. Injections were made at IA −5.5, −5.6 and - 5.7 mm from bregma; ML 0 from ear bars; DV −5.7mm from surface. Mice remained anesthetized for 60 minutes prior to perfusion with 4% paraformaldehyde. The experimenter performing the injection was blinded to group.

### Tissue Processing

Animals were deeply anesthetized (100mg/kg sodium pentobarbital) and transcardially perfused with heparinized saline followed by 4% paraformaldehyde 24 h after the second saline injection. The brain was removed, the medulla blocked and embedded in optimal tissue cutting compound (OCT), cryopreserved in 30% sucrose overnight, and then frozen at −20°C until analysis. Serial sections were cut on a microscope at 20 μm and placed on slides. These sections included the nucleus raphe magnus, the nucleus raphe obscurus and the nucleus raphe pallidus for each animal.

### Immunohistochemistry

Sections from all animals were then immunohistochemically stained for phosphorylated-NR1 (p-NR1) according to previously published procedures [58]. Sections were incubated overnight in the primary antibody, p-NR1 (Millipore Cat# ABN99, RRID:AB_10807298, 1:500 dilution), followed the next day by 1h incubation with biotinylated goat anti-rabbit (Jackson ImmunoResearch Labs Cat# 111-066-144, RRID:AB_2337970, 1:200) and then 1h incubation in streptavidin-Alexa 568 conjugate Alexa 647-conjugate (Thermo Fisher Scientific Cat# S-21374, RRID:AB_2336066, 1:500). It has previously been shown that downregulation of NMDA receptors in the RVM reduces p-NR1 staining showing specificity of staining [14]. A second set of slides was used to examine tryptophan hydroxylase (TPH) staining as a marker for serotonin neurons using the following staining protocol. Sections were incubated overnight in the primary antibody, anti-Tryptophan Hydroxylase/Tyrosine Hydroxylase/Phenylalanine Hydroxylase (Millipore Cat# MAB5278, RRID:AB_2207684, 1:1000), followed the next day by 1h incubation with the secondary IgG Alexa-647 (Jackson ImmunoResearch Labs Cat# 115-605-206, RRID:AB_2338917, 1:500). Removal of the primary antibody eliminated TPH staining in the tissue.

### Data analysis

All images were mapped to Paxinos and Franklin Mouse atlas [46] to define the region for quantification. A template for each Bregma was made to outline the nuclei within the RVM and NRO/NRP to guide capturing of images to allow standardizing areas counted based on location (Figure 1). Low power phase images (4x; light microscope) were collected to identify location of the section on each slide and use as a landmark for taking higher magnification images (Figure 1). The quantification of the RVM included the nucleus raphe magnus and the gigantocellularis pars alpha. The caudal medulla included the nucleus raphe obscurus and the nucleus raphe pallidus.

Sections with dermorphin-488 staining were imaged with confocal microscopy (Confocal Zeiss 710) for dermorphin-488 (argon laser), Fluoro-Gold (diode laser), and p-NR1 (helium laser) or TPH (helium neon laser) immunoreactivity. A 20x power was used for imaging p-NR1-stained sections and a 10x power was used for imaging TPH-labeled sections. Images were built in Image J to form a composite image of all 3 filters. All cells in dermoprhin-488 labeled sections were counted separately for each filter as positively or negatively labeled. All cells were counted by 1 of two individuals for each animal and data were summarized for each individual animal. Both investigators were blinded to group. The two investigators that counted the cells were trained by the same investigator, and interrater reliability of the counts was determined to be r>0.9. We then calculated the percentage of dermorphin-488-positive cells that stained for p-NR1 (or TPH), Fluoro-Gold, and for p-NR1 (or TPH) and Fluoro-Gold.

Statistical analysis was done with a multivariate ANOVA for injury (pH 4, pH 7.2), and exercise condition (sedentary, active). The percent of dermorphin-488-positive cells that expressed Fluoro-Gold, p-NR1 or TPH or Fluoro-Gold+p-NR1 or TPH were analyzed and reported. For comparison, we also calculated the number of Fluoro-Gold-labeled cells that expressed either p-NR1 or TPH. Data are expressed as the mean with S.E.M. for each condition: injury or exercise condition for the rostral medulla and caudal medulla.

## Results

Since ON-cells in the RVM facilitate pain and removal of ON-cells reduces hyperalgesia after nerve injury [33,49], we examined if there were differences in ON-cells in the RVM after chronic pain and after exercise. To label ON-cells, we microinjected the mu-opioid agonist dermorphin-488 into the NRM, NRO and NRP in anesthetized animals.

Fluorescent microscopic imaging shows strong labeling for dermorphin-488 in the RVM and the NRO/NRP around the sites of injection (Figure 2).

**Figure 2.**
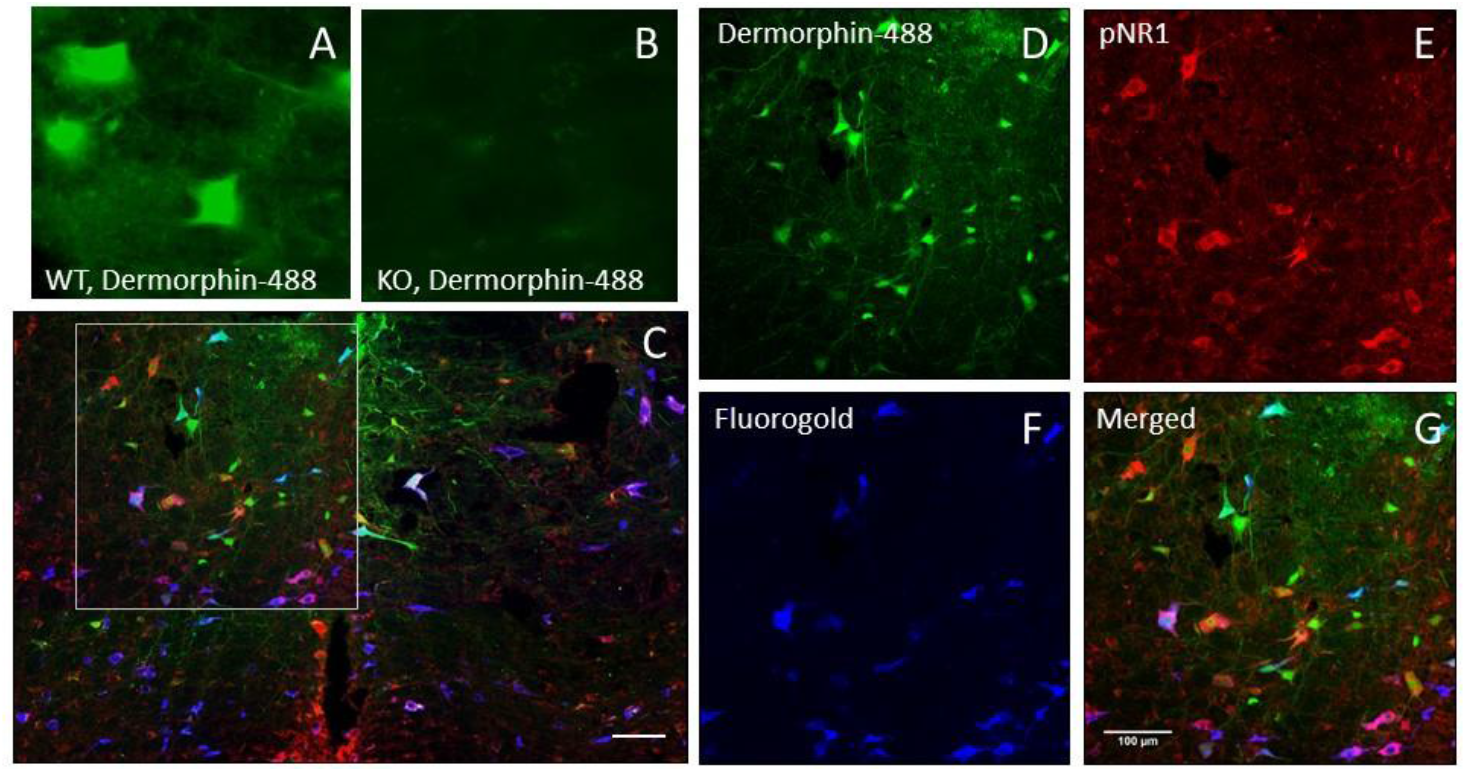
A. Dermorphin-488 immunofluorescence from WT mice. B. Dermorphin-488 fluorescence from mu-opioid receptor knockout mice C. Representative image from the RVM with for dermorphin-488 (green), p-NR1 (red), and Fluoro-Gold (blue). Bar = 50μm D. Higher magnification image from C (box) showing dermorphin-488 immunofluorescence. Bar = 100μm E. Higher magnification image from C (box) showing p-NR1 immunoreactivity. Bar = 100μm. F. Higher magnification image from C (box) showing Fluoro-Gold immunofluorescence. Bar = 100μm. G. Higher magnification of the merged images of D,E.F showing overlap of immunofluorescence.

Since prior studies show alterations in p-NR1 in the NRM, NRO and NRP in a chronic muscle pain model that is modulated by physical activity [58], we immunostained the tissue for p-NR1. Physical activity was induced by provided running wheels to mice; mice averaged 6.1 ± 8.6 km/day. Nearly all dermorphin-positive cells stained for p-NR1. To determine if there were differences in the population of ON-cells that project to the spinal cord, we placed Fluoro-Gold in the dorsal horn of the spinal cord to retrogradely label medullary cells that project to the dorsal horn. A subpopulation of dermorphin-positive cells projected to the spinal cord.

In the p-NR1 staining groups, we counted 282 ± 43 cells per animal for dermorphin-488 and 219 ± 34 cells for Fluoro-Gold in the RVM, and an average of 60 ± 9 cells for dermorphin-488 and 54 ± 7 cells for Fluoro-Gold in the NRO/NRP. Three animals did not have labeled dermorphin-488 cells in the NRO/NRP and were excluded (male runner pH 4.0, female runner pH 4.0, male sedentary pH 4.0). The majority of dermorphin-488-positive cells in the RVM and the NRO/NRP were positively labeled with p-NR1 (Figure 3). A subpopulation of dermorphin-488-positive cells projected to the spinal cord from the RVM (69±11%, mean±SD) in sedentary animals. There was a significant reduction in dermorphin-488-positive cells projecting to the spinal cord from the RVM which decreased to 47 ± 11% (mean+SD) in animals that were physically active. Similar decreases in spinally projecting cells were observed for dermorphin-488-positive cells stained for p-NR1 and labeled with Fluoro-Gold in the RVM. A between-subjects effect for activity status was found for the RVM for the dermoprhin-488+/FG+ group (F1,20=14.4, p=0.001) and the dermorphin-488+/FG+/p-NR1+ group (F1,20=12.7, p=0.002). There were no differences for chronic pain status (pH4, pH 7.2) for any of the analyses. A secondary analysis showed no difference in spinally projecting cells positive for NR1 (FG+/p-NR1+).

**Figure 3.**
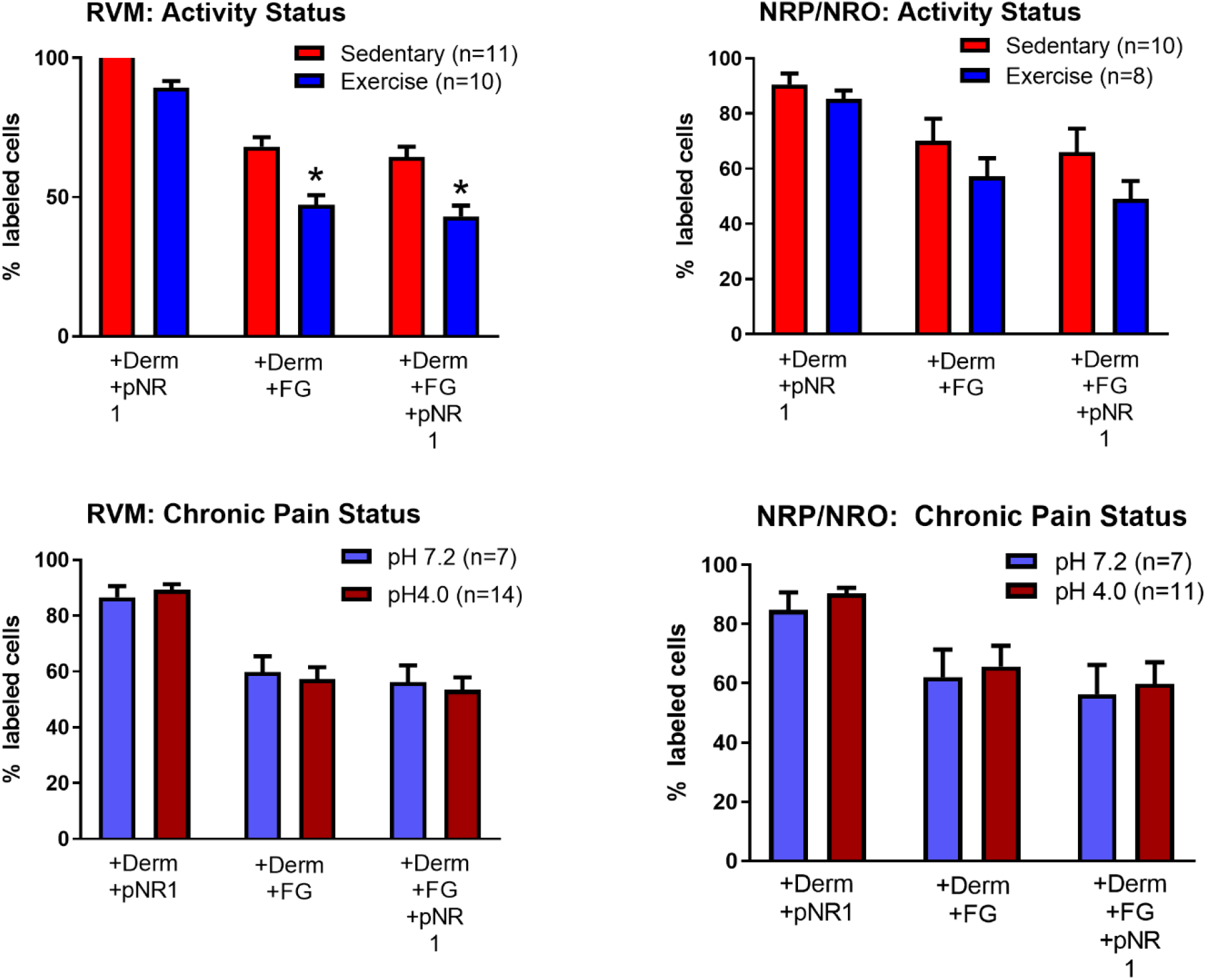
Summary of the percentage of dermorphin-488-positive cells that also stain for p-NR1, Flourogold (FG) or Fluoro-Gold and p-NR1 by different conditions. Nearly all dermorphin-positive cells in the RVM and NRP/NRO stain for p-NR1 and approximately 60% of these cells project to the spinal cord (+FG) in sedentary animals. The proportion of cells projection to the spinal cord is significantly decreased from the RVM, but not the NRP/NRO in animals that were physically active. Data are mean ± SEM

Similar to the RVM, the majority of dermorphin-488-positive cells in the NRO/NRP also stained for p-NR1 and approximately 60% of these cells projected to the spinal cord (Figure 3). However, there was no effect of activity, chronic pain status, or an interaction between activity and chronic pain status in the NRO/NRP (Table 1).

**Table 1.** Statistical analysis for sections labeled with dermorphin-488, p-NR1, and Fluoro-Gold by site: Nucleus Raphe Magnus (NRM) and Nucleus Raphe Obscurus/Nucleus Raphe Pallidus (NRO/NRP) analyzed for activity status (wheel running or sedentary) and pain status (chronic muscle pain, sham), and an interaction between activity and pain status), or sex (male, female).

| **Table 1** | **Site** | **Activity Status** | **Pain Status** | **Activity\*Pain Interaction** |
| --- | --- | --- | --- | --- |
| ***Dermorphin+/p-NR1+*** | NRM | $F_{1,20}=0.001,$<br>$p=0.98$ | $F_{1,20}=.48$<br>$p=0.5$ | $F_{1,20}=.001$<br>$p=0.24$ |
| ***Dermorphin+/FG+*** | NRM | **$F_{1,20}=14.4,$**<br>**$p=0.001$** | $F_{1,20}=0.05,$<br>$p=0.83$ | $F_{1,20}=.04$<br>$p=0.84$ |
| ***Dermorphin+/FG+/p-NR1+*** | NRM | **$F_{1,20}=12.7,$**<br>**$p=0.002$** | $F_{1,20}=0.05,$<br>$p=0.82$ | $F_{1,20}=.006$<br>$p=0.94$ |
| ***Dermorphin+/p-NR1+*** | NRO/NRP | $F_{1,17}=0.60,$<br>$p=0.45$ | $F_{1,17}=0.93$<br>$p=0.35$ | $F_{1,17}=.76$<br>$p=0.4$ |
| ***Dermorphin+/FG+*** | NRO/NRP | $F_{1,17}=.83,$<br>$p=0.38$ | $F_{1,17}=0.05,$<br>$p=0.82$ | $F_{1,17}=.83$<br>$p=0.38$ |
| ***Dermorphin+/FG+/p-NR1+*** | NRO/NRP | $F_{1,17}=0.1.5,$<br>$p=0.24$ | $F_{1,17}=0.05,$<br>$p=0.82$ | $F_{1,20}=.99$<br>$p=0.33$ |

Since RVM and the NRO/NRP are key serotonergic nuclei, and increased serotonin in the RVM can reduce pain [6,39,51], we examined if there were differences in TPH immunoreactivity in dermorphin-488-positive cells. Figure 4 shows representative images of dermorphin-488, TPH immunoreactivity, and Fluoro-Gold in the RVM. In this group we counted an average 107 ± 10 cells positive for TPH, 166 ± 31 cells positive for dermophin-488 and 256 ± 26 cells positive for Fluro-Gold per animal in the RVM, and 22 ± 4 cells positive for TPH, 45 ± 10 cells positive for dermorphin-488 and an average of 36 ± 5 cells positive for Fluoro-Gold in the NRO/NRP. For the RVM, there was minimal co-localization between TPH and dermorphin-488 (≈10%), and between TPH, dermorphin-488 and Fluoro-Gold (≈5%). A greater proportion (≈20-30%) of dermorphin-488-positive cells were labeled for TPH in the NRO/NRP, and ≈10% of these projected to the spinal cord (Figure 5). There were no significant differences between conditions (activity level, chronic pain status) or an interaction between activity level and chronic pain status for TPH positive staining in the RVM (Table 2).

**Table 2.** Statistical analysis for sections labeled with dermorphin-488, TPH (tryptophan hydroxylase), and Fluoro-Gold by site: Nucleus Raphe Magnus (NRM) and Nucleus Raphe Obscurus/Nucleus Raphe Pallidus (NRO/NRP) analyzed for activity status (wheel running or sedentary), pain status (chronic muscle pain, sham), and an interaction between activity and pain status, or sex (male, female).

| **Table 2** | **Site** | **Activity Status** | **Pain Status** | **Activity\*Pain Interaction** |
| --- | --- | --- | --- | --- |
| ***Dermorphin+/TPH+*** | NRM | $F_{1,14}=1.0$,<br>p=0.33 | $F_{1,14}=0.01$<br>p=0.91 | $F_{1,14}=0.39$<br>p=0.54 |
| ***Dermorphin+/FG+/TPH +*** | NRM | $F_{1,14}=3.9$,<br>p=0.08 | $F_{1,14}=0.76$,<br>p=0.4 | $F_{1,13}=0.14$<br>p=0.71 |
| ***Dermorphin+/TPH +*** | NRO/NRP | $F_{1,14}=0.11$,<br>p=0.74 | $F_{1,14}=0.11$<br>p=0.74 | $F_{1,14}=0.001$<br>p=0.97 |
| ***Dermorphin+/FG+/TPH +*** | NRO/NRP | $F_{1,14}=0.12$,<br>p=0.73 | $F_{1,14}=0.12$,<br>p=0.73 | $F_{1,13}=0.31$<br>p=0.79 |

**Figure 4.**
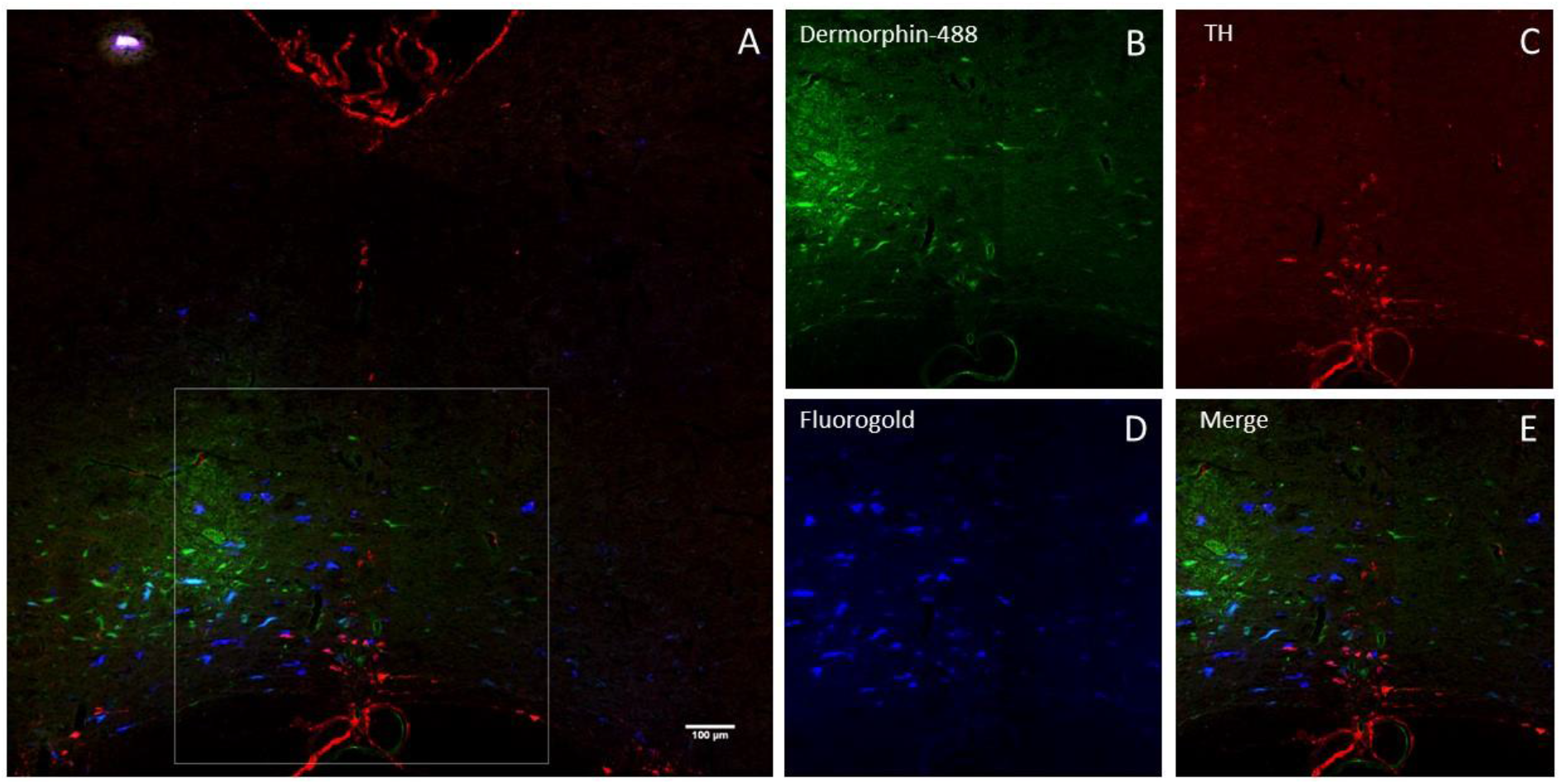
A. Merged image of dermorphin-488, TPH immunofluorescence, and Fluoro-Gold of the RVM. Bar = 50μm. B, C, D, E. Higher power images from A (box) of dermorphin-488 from WT mice (B), TPH (C), Fluro-Gold (D) and the merged image E. Bar = 5μm.

**Figure 5:**
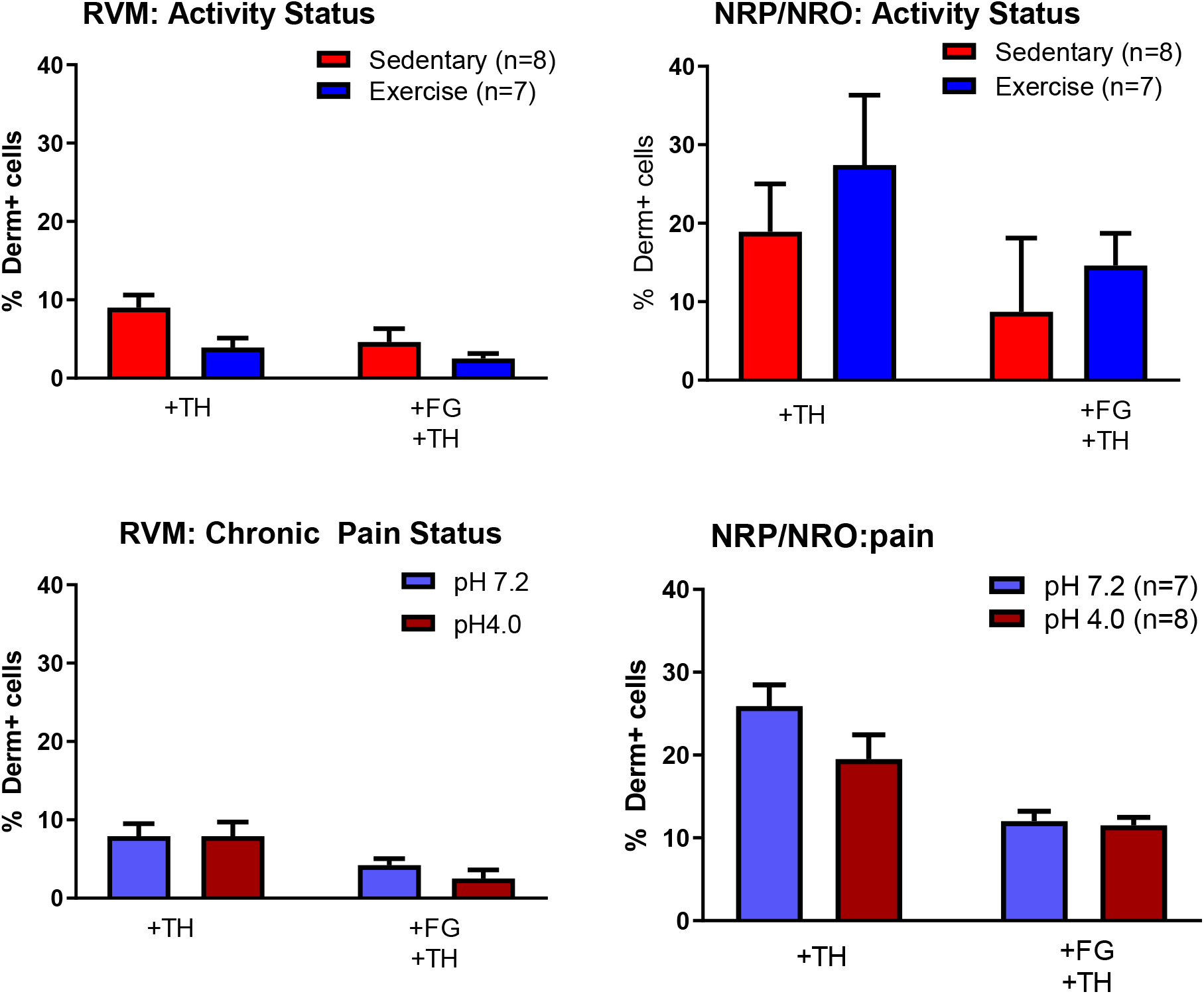
Summary of the percentage of dermorphin-488-positive cells that also stain for TPH or Fluoro-Gold and TPH by different conditions. No significant differences were observed between sedentary and exercise mice, nor for chronic pain status vs controls (Table 1). Data are mean ± SEM.

## Discussion

The current study showed that physically active animals have less dermorphin-488-positive neurons, but not TPH-positive neurons, projecting to the spinal cord when compared to sedentary animals in the RVM, but not the NRO/NRP. These data are consistent with prior studies showing regular exercise produces analgesia through endogenous opioid systems in the RVM and spinal cord [6,39,60]. Since mu-opioid receptors are purported to be ON-cells and facilitate nociception [17,24], these data suggest that there is less descending facilitation from the RVM in physically active mice.

Classically, the RVM is thought to modulate nociception while the NRO/NRP are thought to regulate motor function; however, there is overlap of function between these two regions with some neurons in the NRO/NRP responding to nociceptive input and some in the RVM responding to motor input. Both nuclei express mu-opioid receptors, p-NR1 and contain serotonergic neurons [23,58], alter expression of p-NR1 and the serotonin transporter after chronic muscle pain and exercise [6,39,58], and blockade of NMDA receptors in both nuclei is antinociceptive [13,14,56,62,67,68].

Projections from RVM are found in all laminae in the dorsal and ventral horn [28], and projections from the NRO and NRP are found to project to the deep dorsal horn, intermediate zone, and ventral horn [38,42]. The current study is consistent with these prior findings showing the majority of Fluoro-Gold projections from the dorsal horn to the RVM, but still showing some projections to NRO and NRP. The current study showed approximately 60% mu-opioid positive cells from the RVM projected to the spinal cord in sedentary animals, and approximately 10% of dermorphin-488-positive cells were labeled for TPH agreeing with prior literature [17,41]. We extended these prior studies to include the NRP/NRO and show similar projections. The distinct function between these two nuclei and their role in pain and exercise-induced analgesia will need to be investigated in future studies in more detail.

It is unclear if the reduction in mu-opioid-expressing projections to the spinal cord are due to a functional or structural change in the connections. It is possible that physical activity changes the phenotype of RVM neurons so there are less ON-cells. A change in phenotype in the RVM is supported by prior literature which showed that neutral cells adopted both ON-cell and OFF-cell phenotypes after inflammation resulting in increases in both ON-cells and OFF-cells [43]. Exercise can alter neuron phenotype as evidenced by a prior study showing fast, but not slow or intermediate, motor neurons increase expression of glutamate after exercise training [2].

Alternatively, exercise could enhance synaptogenesis of OFF cell projections in the spinal cord, purported to inhibit nociception, which would result in a lower proportion of ON-cells. In support, there is a substantial body of literature showing cell proliferation, increased synaptic densities, and synaptogenesis in the hippocampus after exercise [64-66]. For example, in the hippocampus, blockade of mu-opioid receptors reduces exercise-induced synaptogenesis suggesting endogenous opioids can promote synaptogenesis [47]. This is in contrast to exogenous opioids, such as morphine, where chronic administration reduces synaptogenesis in the hippocampus [15]. It is unclear if the effects of opioids on synaptogenesis extend to endogenous opioids in the spinal cord and anti-nociceptive pathways.

Finally, exercise could increase the opioid tone in the RVM that resulting in desensitization of mu-opioid receptors. Consistent with this hypothesis, prior studies show that acute blockade of opioid receptors with naloxone in the RVM reverses the analgesia produced by running wheel activity [6] suggesting increased release of opioids and continued activation of opioid receptors. Previous studies show there are increases in p-NR1 in sedentary animals after muscle insult in the RVM and NRO/NRP that did not occur in physically active animals [20,56,58]; however in the current study there were no changes in expression of p-NR1 in the mu-opioid expressing cells. In fact, the majority of mu-opioid receptor expressing cells also expressed p-NR1 regardless of conditions. If there were decreases in the overall expression of mu-opioid receptor expressing cells with exercise, the p-NR1 decreases would not be detectable, consistent with the current data.

The RVM both facilitates and inhibits nociceptive information through descending input to the spinal cord. ON cells facilitate pain and increase their firing in response to noxious stimuli while OFF cells inhibit pain and decrease their firing in response to noxious stimuli. There is generally a balance between inhibition and excitation. Peripheral injury can shift the balance such that ON cell activity on-cell activity outweighs OFF cell activity after tissue injury [7,32,33,68]. Further evidence shows that removal of ON-cells in the RVM with a dermophin-saporin conjugate reduces hyperalgesia after nerve injury and in visceral pain, reduces MOR protein and mRNA, and DAMGO analgesia [11,49,54,70]. These data support the notion that ON-cells facilitation hyperalgesia.

TPH-expressing neurons are reported to be neutral cells [50,51], but spinal serotonin can either facilitate or inhibit nociception [45]. There is substantial evidence suggests that supraspinal serotonergic input to the spinal cord has facilitatory effects on spinal neurons following tissue injury, primarily through its action on 5-HT3 receptors [21,44,54,61]. On the other hand, electrical stimulation of the RVM produces analgesia, releases serotonin and 5-HT-induced nociception is blocked by 5-HT1 antagonists [29,31,53,59]. Further, endogenous analgesia produced by transcutaneous electrical nerve stimulation or joint mobilization is prevented by spinal blockade of 5-HT1 or 5-HT2 receptor antagonists [52,55]. It has also previously been shown that there are increases in expression of the serotonin transporter in the RVM and NRO/NRP after muscle insult [6,39]; however the current study does not show changes in TPH expression after muscle insult or regular exercise. Prior studies, using electrophysiology show a functional change in phenotype of RVM neurons. Neutral cells, thought to be TPH+ cells, developed ON-cell or OFF-cell like properties after peripheral inflammation [43] while ON-cells and OFF-cells develop new responsiveness to innocuous mechanical stimuli after nerve injury [12]. The current study, using an anatomical approach, suggests that muscle insult does not change phenotype of ON-cells (MOR+) or neutral-cells (TPH+), despite playing a significant role in the development and maintenance of chronic muscle pain [13,14,63].

There are several limitations with the current study. We did not use stereological analysis of labeled cells, but rather counted profiles. We recognize counting profiles could result in an overrepresentation of larger cells and underrepresentation of smaller cells[36]. Another limitation is our inability to determine if the number of mu-opioid receptors were changed in these sections since we used a direct injection of dermorphin-488 to label mu-opioid positive neurons. This was necessary since we were unable to obtain a specific antibody that showed adequate staining in a wild-type mouse compared to a mu-opioid receptor mouse.

In summary, we show that regular exercise reduces the number of putative ON-cells cells that project to the spinal cord. These data suggest that regular exercise alters central facilitation so that there is less descending facilitation to result in a net increase in inhibition. This change in the balance between inhibition and facilitation could explain why regular exercise is protective against the development chronic pain.

